# The landscape of myeloid and astrocyte phenotypes in acute multiple sclerosis lesions

**DOI:** 10.1101/632554

**Authors:** Calvin Park, Gerald Ponath, Maya Levine-Ritterman, Edward Bull, Eric C. Swanson, Philip L. De Jager, Benjamin M. Segal, David Pitt

## Abstract

Activated myeloid cells and astrocytes are the predominant cell types in active multiple sclerosis (MS) lesions. Both cell types can adopt diverse functional states that play critical roles in lesion formation and resolution. In order to identify phenotypic subsets of myeloid cells and astrocytes, we profiled acute MS lesions with thirteen glial activation markers using imaging mass cytometry (IMC), a method for multiplexed labeling of histological sections. In a demyelinating lesion, we found multiple distinct myeloid and astrocyte phenotypes that populated separate lesion zones. In a post-demyelinating lesion, phenotypes were less distinct and more uniformly distributed. In both lesions cell-to-cell interactions were not random, but occurred between specific glial subpopulations and lymphocytes. Finally, we demonstrated that myeloid, but not astrocyte phenotypes were activated along a lesion rim-to-center gradient, and that marker expression in glial cells at the lesion rim was driven more by cell-extrinsic factors than in cells at the center. This proof-of-concept study demonstrates that highly multiplexed tissue imaging, combined with the appropriate computational tools, is a powerful approach to study heterogeneity, spatial distribution and cellular interactions in the context of MS lesions. Identifying glial phenotypes and their interactions at different lesion stages may provide novel therapeutic targets for inhibiting acute demyelination and low-grade, chronic inflammation.

## Introduction

Multiple sclerosis (MS) is a common neurological disease, characterized by formation of inflammatory demyelinating lesions in the central nervous system (CNS) [27]. Inflammation is driven by infiltrating lymphocytes and monocytes, in concert with resident activated microglia and astrocytes. Macrophages and reactive astrocytes are the most abundant cell types in acute lesions [19,31]. These cells are highly plastic and can adopt pro-inflammatory, anti-inflammatory, neurotoxic, neuroprotective, and tissue-regenerating functions [4,6,7,21,24,30,31,43].

Previous studies have identified macrophage phenotypes in MS lesions based on the expression of single classical (M1) and alternative (M2) activation markers; however, those studies have produced limited and sometimes conflicting results [6,43]. A recent study of traumatic brain injury in mice demonstrated random expression of M1/M2 polarization markers in infiltrating macrophages in seemingly incompatible combinations [17]. This suggests that the M1/M2 polarization paradigm, which originated as an *in vitro* concept, is of limited value for distinguishing myeloid phenotypes in the inflamed CNS [36].

Current studies, including our own, are using single-cell or single-nuclear RNA sequencing to comprehensively assess the complex phenotypes of human glial cells in healthy and diseased brains [14,23,28]. Through these efforts, we and others have observed varying levels of myeloid cell/microglial diversity in the human brain depending on the neurological disease and clustering aims, where diverse functional states of microglia are associated to varying degrees with different brain pathologies. Moreover, since RNA sequencing techniques do not capture spatial information, the distribution and interactions of these subpopulations in CNS tissue have not yet been examined.

Several novel techniques make it now possible to perform high parameter imaging of histological section and evaluate complex cellular phenotypes *in situ* [5,8,9,11,42]. Imaging mass cytometry (IMC), like mass cytometry (CyTOF), relies on metal isotope-labeled antibodies and combines immunohistochemistry with high-resolution laser ablation followed by time-of-flight mass spectrometry [9,44]. This approach allows for simultaneous quantitative profiling with up to 37 antibodies on a single tissue section at subcellular resolution. Moreover, novel computational tools have become available to extract single-cell information from highly multiplexed histological data [3,25,37].

In this proof-of-concept study, we applied IMC and single-cell analytics to two active MS lesions – one demyelinating and one post-demyelinating – to examine the cellular heterogeneity of myeloid cells and astrocytes, based on thirteen markers that are known to be expressed by activated glial cells in MS lesions [2,6,13,15,32,43,45,46]. We demonstrate that multiplexed tissue imaging, in combination with the appropriate computational tools, can extract previously unattainable information from histological sections, including definition of cellular subpopulations, their distribution within the lesion environment, specific cell-cell interactions, phenotypic transitions and the impact of spatial sources on marker expression.

## Materials and methods

### MS lesions

Human CNS tissue was obtained at autopsy from two patients with relapsing-remitting MS according to Institutional Review Board-approved protocols. After autopsy, brain tissue was fixed in 10% formalin, and lesions were cut based on MRI. Lesion tissue was subsequently embedded in paraffin and sectioned at 5 μm thickness.

A highly inflamed, active lesion was chosen for analysis from each patient: the demyelinating lesion was selected from a 42-year-old male with 5 years disease duration (5.5 hour *post mortem* interval), while the post-demyelinating lesion was chosen from a 32-year-old female with 6 years disease duration (8 hour *post mortem* interval). Lesions from both patients have been characterized in previous studies [10,29].

### Brightfield histology

For basic characterization, lesions were stained against CD68, myelin basic protein (MBP) and neuronal marker MAP2, and examined with brightfield microscopy. De-paraffinized and rehydrated sections were subjected to antigen retrieval in pH 6, 10 mM citrate buffer at 96°C for 20 minutes, cooled, quenched in 0.3% peroxide, and blocked with FC receptor binding inhibitor and normal serum before incubation with primary antibody (Additional file 1: Table S1) overnight at 4°C. Sections were subsequently incubated with appropriate biotinylated secondary antibodies, processed with an avidin/biotin staining kit with 3,3-diaminobenzidene (DAB) as the chromogen (Vector ABC Elite Kit and DAB Kit, Vector Laboratories), then counterstained with hematoxylin [29]. Adequate controls using isotype control antibodies were performed for each primary antibody. Sections were rinsed with distilled water, dehydrated, and cover-slipped with Permount (Vector Laboratories). Images were acquired using a Leica DM5000 B microscope with a Leica color camera DFC310 Fx and Leica Application Suite (version 4.9.0) imaging software. Images were processed with Panoramic Viewer (3DHISTECH) and Photoshop (Adobe) software.

### Antibody validation and conjugation to metal isotopes for IMC

Several lanthanide-conjugated antibodies were purchased from Fluidigm. Antibodies not available in metal-conjugated form were purchased in carrier-free solution and validated by brightfield immunohistochemistry using the appropriate isotype control antibodies. Subsequently, antibodies were conjugated to lanthanide metal isotopes following the Maxpar® Antibody Labeling Kit protocol (Fluidigm). Metal-conjugated antibodies were stored at 0.5 mg/mL in PBS-based Antibody Stabilizer (Candor Bioscience) with 0.05% sodium azide at 4°C. Working concentrations for all metal-conjugated antibodies were optimized by IMC (Additional file 1: Table S2) on MS lesion tissue.

### Imaging mass cytometry

For IMC histology, tissue sections were de-paraffinized and rehydrated, and antigen retrieval was performed in pH 8, 1 mM EDTA buffer at 96°C for 20 minutes. Sections were cooled at room temperature and rinsed in tap water and TBS (20 mM Tris with 500 mM NaCl, pH 7.5). Tissue was blocked for 3 hours at room temperature with 0.3% BSA, 25% FBS and 0.1 mg/mL FC receptor binding inhibitor in TBS-T (TBS + 0.05% Tween-20). All antibodies (Additional file 1: Table S2) were diluted in 0.3% BSA in TBS-T and applied to the tissue for overnight incubation at 4°C. Sections were then rinsed in TBS-T and TBS, and counterstained with 125 nM Maxpar® Intercalator-Ir (Fluidigm) in PBS for 1 hour at room temperature. Sections were rinsed in TBS-T, TBS and two washes of distilled water before air-drying at room temperature. Antibody-labeled tissue areas (1000×1000 μm) were raster-ablated using a Hyperion^™^ Laser Scanning Module (Fluidigm) with a 1 μm diameter spot size at 200 Hz. This process was coupled to a Helios^™^ Mass Cytometer (Fluidigm) for lanthanide metal detection [44]. Images for each antibody channel were acquired on CyTOF Software (Fluidigm, version 6.7). MCD Viewer (Fluidigm, version 1.0) was used to export raw 16-bit tiff images for computational analyses on histoCAT (version 1.75) [37]. For visualization purposes, images were processed in MCD Viewer and ImageJ [39].

### Computational analyses

#### Single-cell segmentation

Merged CD68 (macrophages/microglia), S100B (astrocytes), and CD3 (T cells) antibody channel images were used to segment single cell objects on CellProfiler (version 3.0.0) [16]. The resulting segmentation mask images, which outline cell borders, were loaded into histoCAT with corresponding antibody channel images.

#### Identification of cellular phenotypes

Mean single-cell marker intensity values were extracted from segmented antibody channel images on histoCAT and Z-score normalized. Based on the expression intensities of thirteen markers (Additional file 1: Table S2), cell clusters were defined using the PhenoGraph algorithm [20] integrated into histoCAT. Default parameters with 75 nearest neighbors for the early lesion and 50 nearest neighbors for the late lesion were used. These nearest neighbor values were chosen such that over- and under-clustering of phenotypes were avoided. Additional normalization steps were performed internally, as previously described [37].

#### Analysis of cellular phenotypes

To visualize clusters, the Barnes-Hut t-SNE algorithm implemented in histoCAT was executed with the same image and marker inputs used in PhenoGraph, as well as default parameters (initial dimensions, 110; perplexity, 30; theta, 0.5) and internal normalization [1,37]. t-SNE plots were colored to highlight cell clusters or lesion samples, or to show relative marker expression intensity. Images of cell phenotypes visualized in the tissue, as well as segmentation masks overlaid with histology images, were generated in histoCAT. For the remaining analyses, “.csv” files containing single-cell parameters were exported from histoCAT and appropriately processed for their application. To produce an expression heatmap for clusters, normalized marker intensity values were processed using the R *ComplexHeatmap* package, which hierarchically clusters single cells within clusters using Ward’s method [38]. Violin plots showing single-cell marker expression variability for each cluster were generated using the R *ggplot2* package [12]. To study phenotype transitions, Potential of Heat-diffusion Affinity-based Transition Embedding (PHATE) mapping and Monocle Pseudotime analyses were performed [25,34,35,41]. PHATE mapping was performed using normalized marker intensity values, with an adjusted number of nearest neighbors (k = 100) and other parameters left as their default specifications. Pseudotime analysis was performed with negative binomial normalization of marker intensity values and q < 1 to incorporate all markers, based on a test for differential expression among phenotypes. Figures showing phenotype cell size and relative quantity of cells per cluster, as well as correlation matrices and corresponding marker scatter plots with linear regression lines were produced in Prism (version 7). FlowJo software (version 10.5.3) was used to visualize single-cell marker data on flow cytometry plots. Images and figures were recolored when necessary in Photoshop.

#### Analysis of cell spatial relationships

To study the spatial relationships of cell clusters, neighborhood analysis was performed on histoCAT using the PhenoGraph-generated clusters pertaining to each lesion. Significant pairwise phenotype interactions and avoidances were determined by an unbiased permutation test which compared the frequency of one cell type neighboring another to the frequency in 999 random permutations of cell cluster labels. Neighbors were identified within 4 μm from each cell, and neighboring frequency was normalized to the number of interacting cells [37]. Analyses were performed for different degrees of significance (p < 0.05 and p < 0.01), and the results were reported on a heatmap. To identify sources of single-cell marker variation, spatial variance component analysis (SVCA) [3] was performed on Python for different lesion zones, using standardized marker intensity values. SVCA plots were generated in R and Prism.

### Statistical analyses

In plots showing phenotype cell size, data represent mean cell areas + standard deviation. Comparisons of phenotype cell sizes were analyzed by one-way ANOVA followed by the Tukey-Kramer multiple comparison test. Comparisons of two samples were carried out by unpaired Student’s t-tests. For correlation analyses, Pearson correlation coefficients were computed. *p < 0.0001.

## Results

### Histology and cell clustering overview

We analyzed one demyelinating and one post-demyelinating MS lesion according to a classification by Kuhlmann et al. [18], referred to throughout this report as the “early” and “late” lesion, respectively. Both lesions were located in the brain stem and characterized by complete loss of myelin, hypercellularity with the highest cellular density at the lesion rim, and diffuse infiltration with foamy macrophages. The laser-scanned area of the early lesion involved predominantly white matter (WM) but also interspersed gray matter (G/WM), while the scanned area of the late lesion consisted of only WM (Fig. 1).

**Fig. 1.**
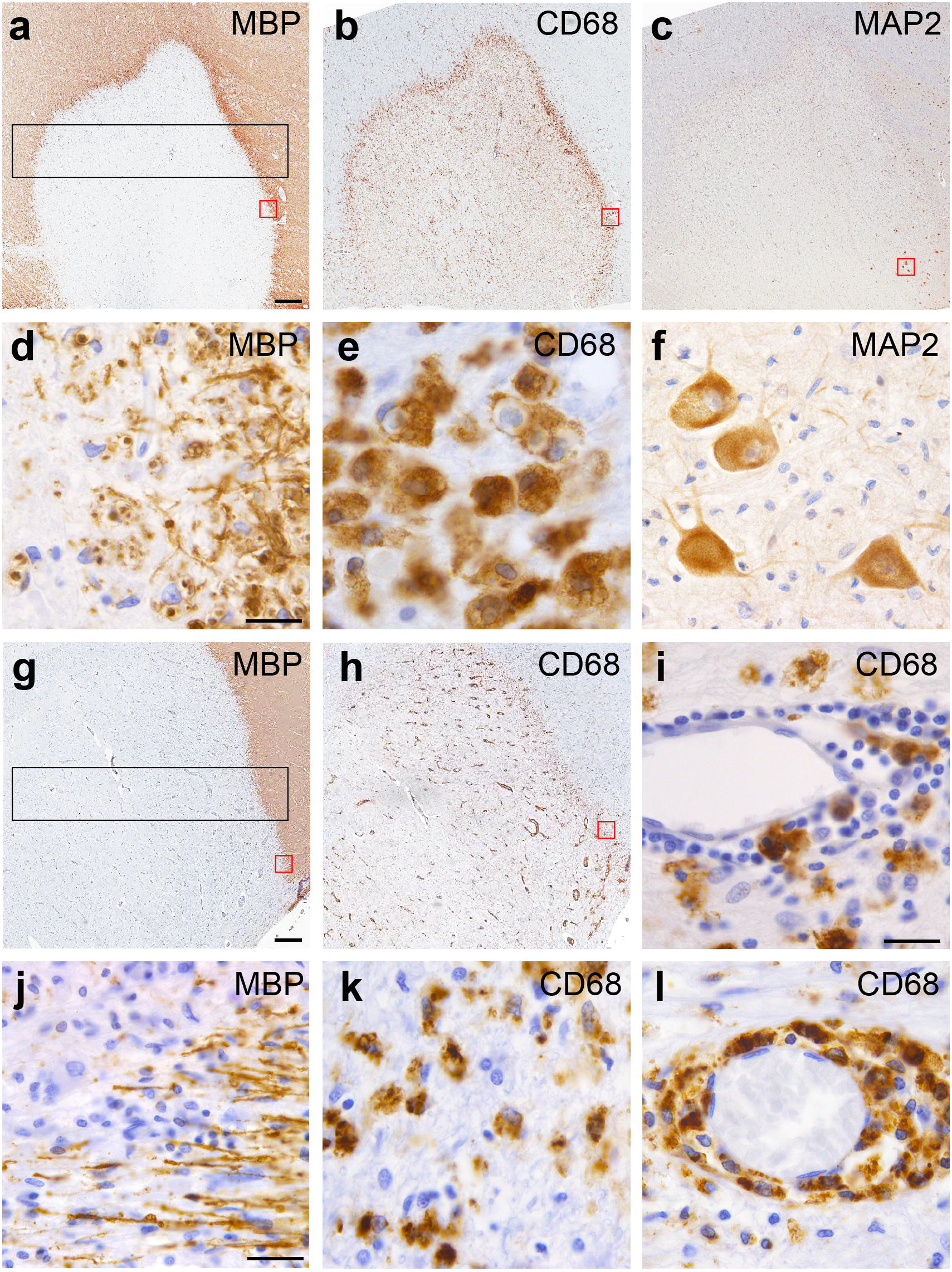
Brightfield images of the MS lesions examined with IMC. **(a-f, i)** Staining of the early, actively demyelinating lesion for **(a)** myelin basic protein (MBP), **(b)** CD68 (macrophages/microglia) and **(c)** the neuronal marker MAP2. The black rectangle in **(a)** shows the region that was laser-ablated for IMC, capturing the NAWM, hypercellular rim on both sides and lesion center. Red boxes in **a-c** correspond to magnified rim areas in **d-f,** demonstrating in **(d)** condensed myelin fragments, indicative of ongoing demyelination, **(e)** dense infiltration with foamy macrophages and **(f)** dispersed neuronal cell bodies in the right lesion rim. **(g, h, j-l)** Staining against MBP and CD68 in the late, post-demyelinating lesion. The black rectangle in **(g)** shows the region laser-ablated for IMC. **(j, k)** Details from **(g)** and **(h)** showing absence of myelin fragments and reduced myeloid cell density at the lesion rim compared to **(e)**. CD68^+^ cells in the perivascular spaces of **(i)** the early lesion and **(l)** the late lesion. Images show hematoxylin counterstaining. Scale bars **a-c, g, h**= 500 μm. Scale bars **d-f, i, j-l** = 25 μm

Consistent with demyelinating activity, macrophages at the rim of the early lesion contained myelin basic protein (MBP)-positive myelin debris, which was absent in macrophages from the late lesion (Fig. 1d, j) [18]. Moreover, foamy macrophages were numerous and large in size in the rim of the early lesion, whereas macrophages in the rim of the late lesion were smaller and less concentrated (Fig. 1e, k).

Perivascular infiltrates in the early lesion contained mostly lymphocytes and only a few undifferentiated monocytes, while the perivascular cuffs in the late lesion consisted predominantly of lipid-laden macrophages, as described previously (Fig. 1i, l) [22,40]. We defined single cells by segmenting cell bodies outlined by the markers CD68 (macrophages/microglia), S100B (astrocytes), and CD3 (T cells) using CellProfiler (Additional file 1: Figure S1) [16]. This allowed for a clear delineation of cell types in the vast majority of cells. In a small fraction of cells, we found overlap between cellular markers, especially between myelin fibers and juxtaposed microglial processes in normal-appearing white matter (NAWM), and between astrocyte processes and closely neighboring macrophages (Additional file 1: Figure S1c, d). We did not exclude these cells in order to avoid bias in our spatial analysis. However, we excluded perivascular lymphocytes from the early lesion, as they were too densely packed to be identifiable as individual cells.

Segmentation masks were overlaid with corresponding images from thirteen antibody channels (Additional file 1: Figures S2, S3) in histoCAT [37], which clustered myeloid cells and astrocytes into phenotypic subpopulations based on their marker expression profiles. The optimal number of phenotype clusters was determined based on separation of the principal cell types (i.e. myeloid cells, astrocytes and T cells) and distinctive expression profiles of phenotypes, using expression heatmaps and t-SNE plots. This resulted in a total of twelve phenotypes for each lesion (Figs. 2, 4).

**Fig. 2.**
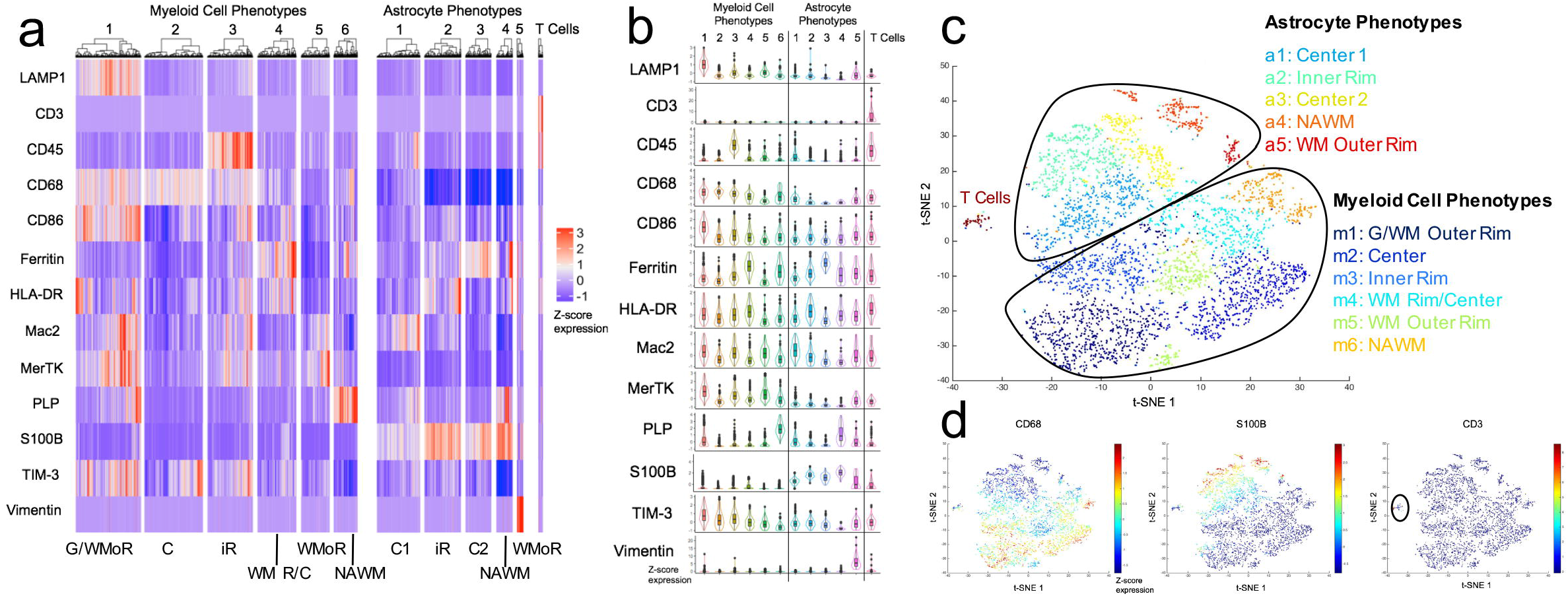
Early lesion cell phenotype profiles. **(a)** Marker expression heatmap for the myeloid, astrocyte and T cell phenotypes, identified by PhenoGraph clustering on histoCAT using segmented cells (n = 4397). The heatmap displays relative expression levels based on Z-score normalized marker intensity values, and single cells are hierarchically clustered within each phenotype group. Labels at the bottom of the heatmap indicate the area of the lesion to which each phenotype localizes. **(b)** Violin plot representation of the data in **(a)**. **(c)** t-SNE plot showing the distinct phenotype clusters. **(d)** t-SNE plot colored by marker intensity, confirming the separation of CD68^+^, S100B^+^ and CD3^+^ cell types. G/WMoR = gray and white matter outer rim; WMoR = white matter outer rim; iR = inner rim; WM R/C = white matter rim/center; C = center; NAWM = normal-appearing white matter

### Phenotype heterogeneity and distribution in early demyelination

In the early lesion, we analyzed a total of 4,397 cells, of which 66.3% were myeloid cells and 32.5% were astrocytes (Additional file 1: Figure S4a). This ratio was higher at the rim than the lesion center. The cells clustered into six myeloid and five astrocyte subtypes (Fig. 2). The myeloid phenotypes were spatially sequestered in four lesional regions, i.e. NAWM, outer lesion rim, inner lesion rim, and center. Within the outer lesion rim, we observed one myeloid cell phenotype at the G/WM interface (m1) and two phenotypes located in the WM rim (m4, m5). The other lesion areas were all occupied by one myeloid phenotype (Fig. 3a). Most myeloid cell activation markers were highly expressed in the outer lesion rim and decreased in intensity towards the lesion center (Fig. 2a, b). The m1 phenotype at the G/WM interface showed the highest activation profile. m5 cells in the outer WM rim were the largest in size, whereas m2 cells in the lesion center were the smallest (Additional file 1: Figure S4b); this may reflect the amount of myelin fragments phagocytosed by foamy macrophages at the advancing lesion edge, and the degradation of myelin in phagocytes within the lesion center [33].

**Fig. 3.**
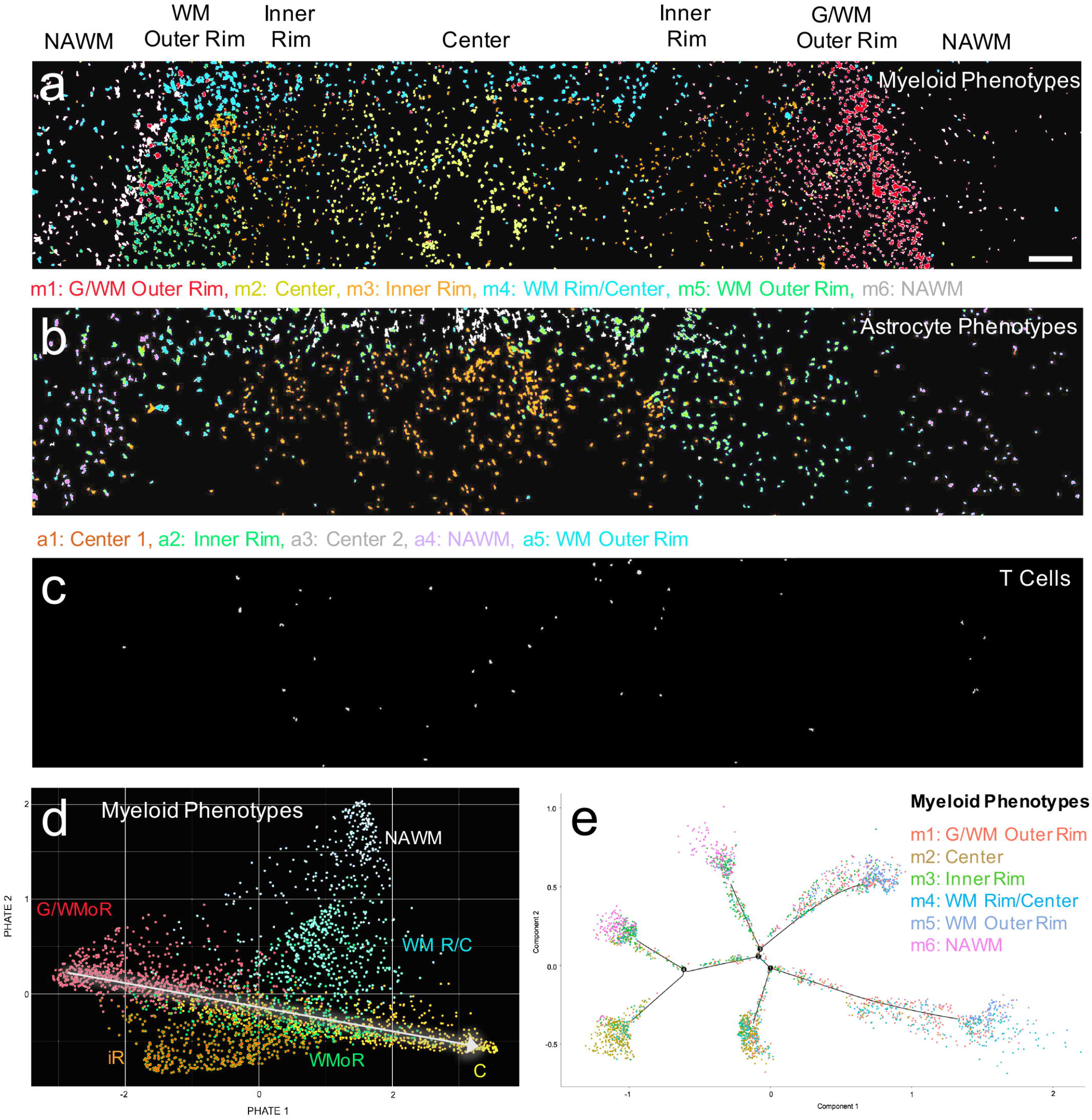
Early lesion phenotype spatial distributions and transition analyses. **(a, b)** Spatial separation of **(a)** myeloid cell and **(b)** astrocyte phenotypes into NAWM, rim and center lesion zones. **(c)** T cells are primarily located in the lesion center. **(d)** PHATE mapping of myeloid cells, indicating that the G/WM outer rim (m1) and lesion center phenotypes (m2) are on a transition continuum (white arrow). **(e)** Pseudotime analysis of myeloid cells shows that phenotypes transition along independent trajectories. Phenotype color schemes on the PHATE and Pseudotime plots reflect the color palettes specific to each analysis. Scale bar for **a-c** = 200 μm. G/WMoR = gray and white matter outer rim; WMoR = white matter outer rim; iR = inner rim; WM R/C = white matter rim/center; C = center; NAWM = normal-appearing white matter

**Fig. 4.**
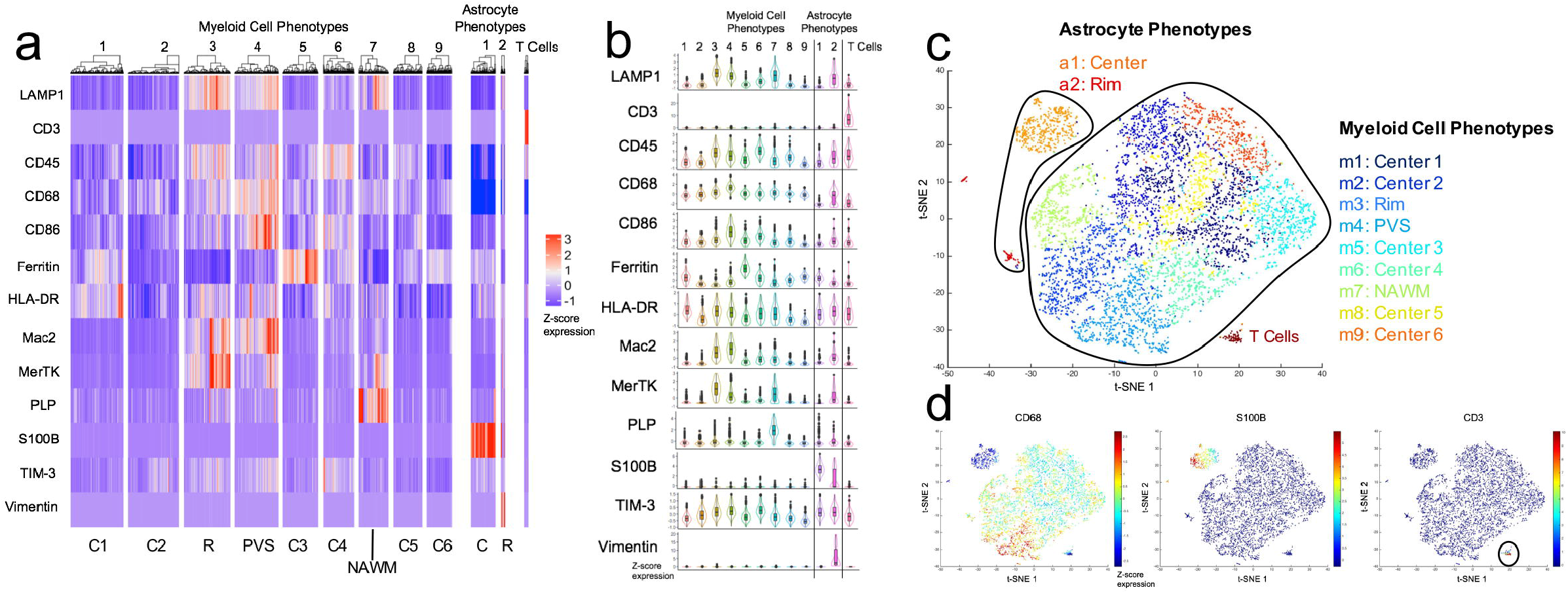
Late lesion cell phenotype profiles. **(a)** Marker expression heatmap for the myeloid, astrocyte and T cell phenotypes, identified by PhenoGraph clustering on histoCAT using segmented cells (n = 6698). The heatmap displays relative expression levels based on Z-score normalized marker intensity values, and single cells are hierarchically clustered within each phenotype group. Labels at the bottom of the heatmap indicate the area of the lesion to which each phenotype localizes. **(b)** Violin plot representation of the data in **(a)**. **(c)** t-SNE plot showing the phenotype clusters. Compared to the early lesion, myeloid cell phenotypes show a low degree of separation. **(d)** t-SNE plot colored by marker intensity, confirming the separation of CD68^+^, S100B^+^ and CD3^+^ cell types. R = rim; C = center; PVS = perivascular space; NAWM = normal-appearing white matter

Astrocyte phenotypes were each defined by one distinct, highly expressed marker (Fig. 2a, b). Analogous to the myeloid phenotypes, the five astrocyte phenotypes were also spatially stratified into the NAWM, lesion rim, and lesion center (Fig. 3b). Furthermore, a5 astrocytes located in the outer rim within WM were larger than all other phenotypes (Additional file 1: Figure S4b). Unlike in myeloid cells, marker expression in astrocytes did not follow a gradient from the rim to the lesion center, but was uniform throughout the lesion.

T cells made up the smallest population of all immune infiltrates, and were concentrated within the lesion center (Fig. 3c; Additional file 1: Figure S4a). These cells uniformly expressed CD45 and HLA-DR (Fig. 2a, b), and did not separate into different clusters.

To determine possible transitions between phenotypes, we applied Potential of Heat-diffusion Affinity-based Transition Embedding (PHATE) mapping and Pseudotime to the myeloid cell and astrocyte populations. PHATE mapping improves on t-SNE by better visualizing phenotype transitions, where smooth continua from one phenotype to another indicate a transition trajectory [25]. Rather than a rim-to-center continuum that aligned with the sequential organization of phenotypes in the lesion, our analysis showed only a partial continuum between the G/WM outer rim phenotype (m1) and the lesion center phenotype (m2) (Fig. 3d), without including the inner rim phenotype (m3). Similarly, analysis with Pseudotime, an alternative tool for visualizing phenotype transitions [41], showed that myeloid phenotypes did not follow a linear transition trajectory but branched into several independent fates (Fig. 3e). PHATE mapping and Pseudotime analysis of astrocyte phenotypes also suggested independent phenotypic fates rather than linear phenotype trajectories (Additional file 1: Figure S5a, Pseudotime analysis not shown).

### Low phenotype heterogeneity and random phenotype distributions in the late demyelinating lesion

In the late, post-demyelinating lesion we analyzed 6,698 cells, with myeloid cells far outnumbering astrocytes (91.1% myeloid cells; Additional file 1: Figure S6a), particularly at the lesion rim. The same clustering criteria used for the early lesion resulted in nine myeloid phenotypes and two astrocyte phenotypes in this late lesion (Fig. 4). Myeloid phenotypes separated into lesion rim (m3), perivascular space (m4), and NAWM (m7) zones (Fig. 5a). In contrast to the early lesion, the other six myeloid phenotypes were intermixed throughout the lesion center. These phenotypes showed a low degree of separation on the t-SNE plot, indicating similar marker expression profiles (Fig. 4c). The phenotypes in the lesion rim and the perivascular space (m3, m4) were characterized by high expression of the majority of markers, and shared a similar expression profile with the G/WM rim phenotype in the early lesion (m1) (Fig. 4a, b). As in the early lesion, myeloid phenotypes in the lesion rim (m3) and perivascular space (m4) were significantly larger than those in the lesion center (Additional file 1: Figure S6b), but were overall smaller than in the early lesion (data not shown).

**Fig. 5.**
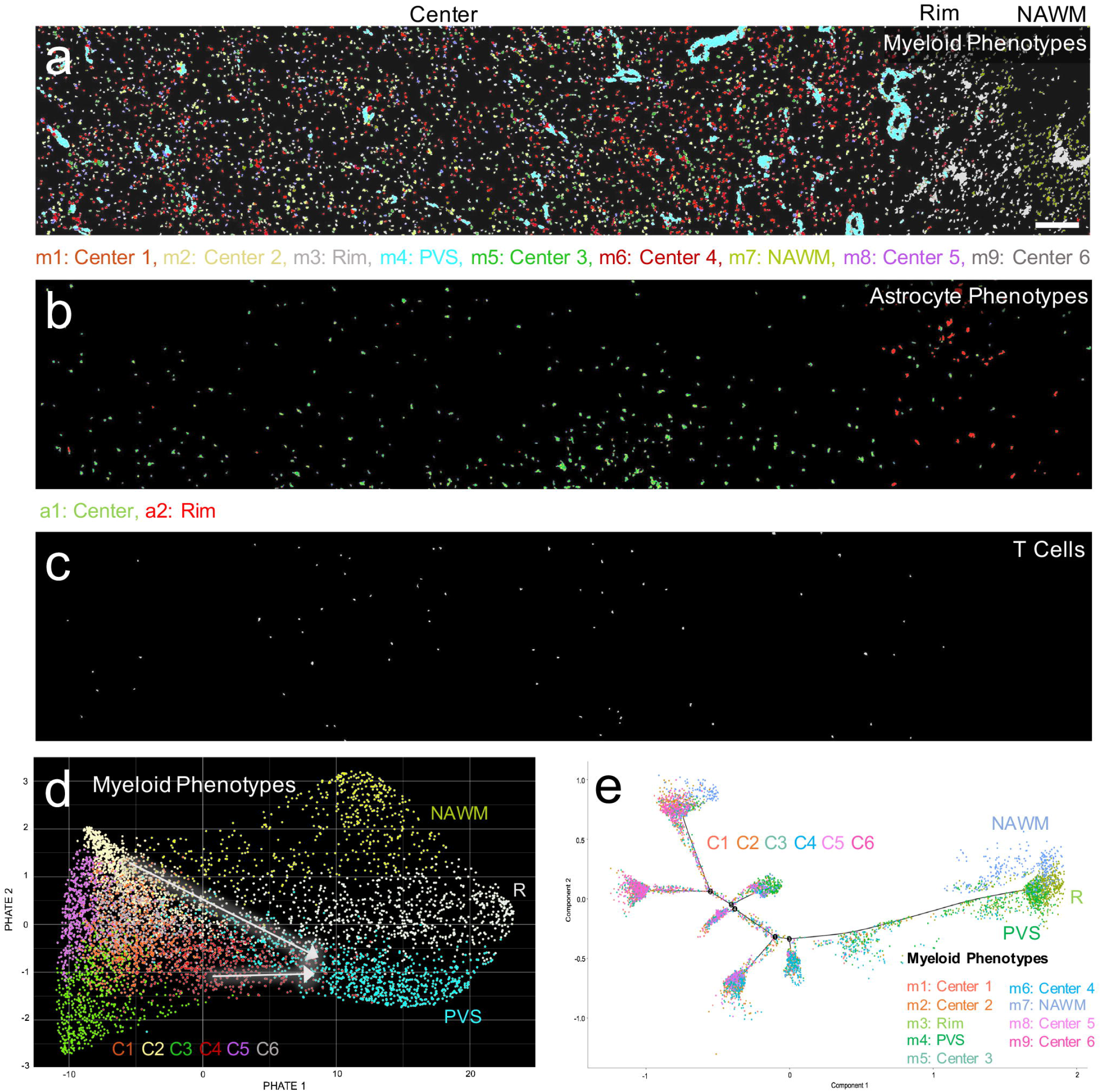
Late lesion phenotype spatial distributions and transition analyses. **(a)** Spatial organization of myeloid cell phenotypes in the lesion. Phenotypes in the rim (m3), perivascular space (m4) and NAWM (m7) separate into distinct zones, while lesion center phenotypes (m1, m2, m5, m6, m8, m9) are uniformly distributed. **(b)** Spatial distribution of astrocyte phenotypes. One phenotype (a1) predominantly occupies the lesion center and the other (a2) occupies the rim. **(c)** T cells are primarily distributed in the lesion center. **(d)** PHATE mapping of myeloid cells, showing that two lesion center phenotypes (m2, m6) are on a continuum with perivascular space cells (m4, white arrows). **(e)** Pseudotime analysis of myeloid cells shows a similar trajectory to PHATE mapping. Phenotype color schemes on the PHATE and Pseudotime plots reflect the color palettes specific to each analysis. Scale bar for **a-c**= 200 μm. R = rim; C = center; PVS = perivascular space; NAWM = normal-appearing white matter

Astrocytes were clustered into two phenotypes, with one phenotype localizing primarily to the lesion rim, and the other to the lesion center (Fig. 5b). The rim phenotype (a2) exhibited a marker expression profile similar to the rim phenotype in the early lesion (a5), (Fig. 4a, b). Finally, T cells were few (Fig. 5c; Additional file 1: Figure S6a), and expressed the activation markers CD45 and HLA-DR (Fig. 4a, b), as observed in the early lesion. To directly compare cell populations in both lesions, we mapped cells from both lesions on the same t-SNE plot. Cell populations overlapped only moderately, highlighting the differences between the phenotypes from each lesion (Additional file 1: Figure S7).

PHATE mapping demonstrated a partial continuum between two lesion center myeloid phenotypes (m2, m6) and the perivascular space phenotype (m4) (Fig. 5d), which was confirmed by Pseudotime (Fig. 5e). These analyses support a lesion center-to-perivascular transition, but not a continuum where all phenotypes align along a rim-to-center transition axis. The same analyses for astrocytes showed substantial overlap between both phenotypes but not a linear transition (Additional file 1: Figure S5b, Pseudotime analysis not shown).

In the early lesion, we found no correlation between the expression intensities of different markers at a single-cell level (Additional file 1: Figure S8a, b). This was true for an analysis of all cells and cells of each phenotype (data not shown). In the late lesion, we found strong correlations between the M2 markers MerTK and Mac2, and MerTK and LAMP1 in both myeloid cells and astrocytes (Additional file 1: Figure S8c, d). This resulted from high and continuous dynamic ranges of marker expression (Fig. 6). Correlations were weak between the other marker pairs.

**Fig. 6.**
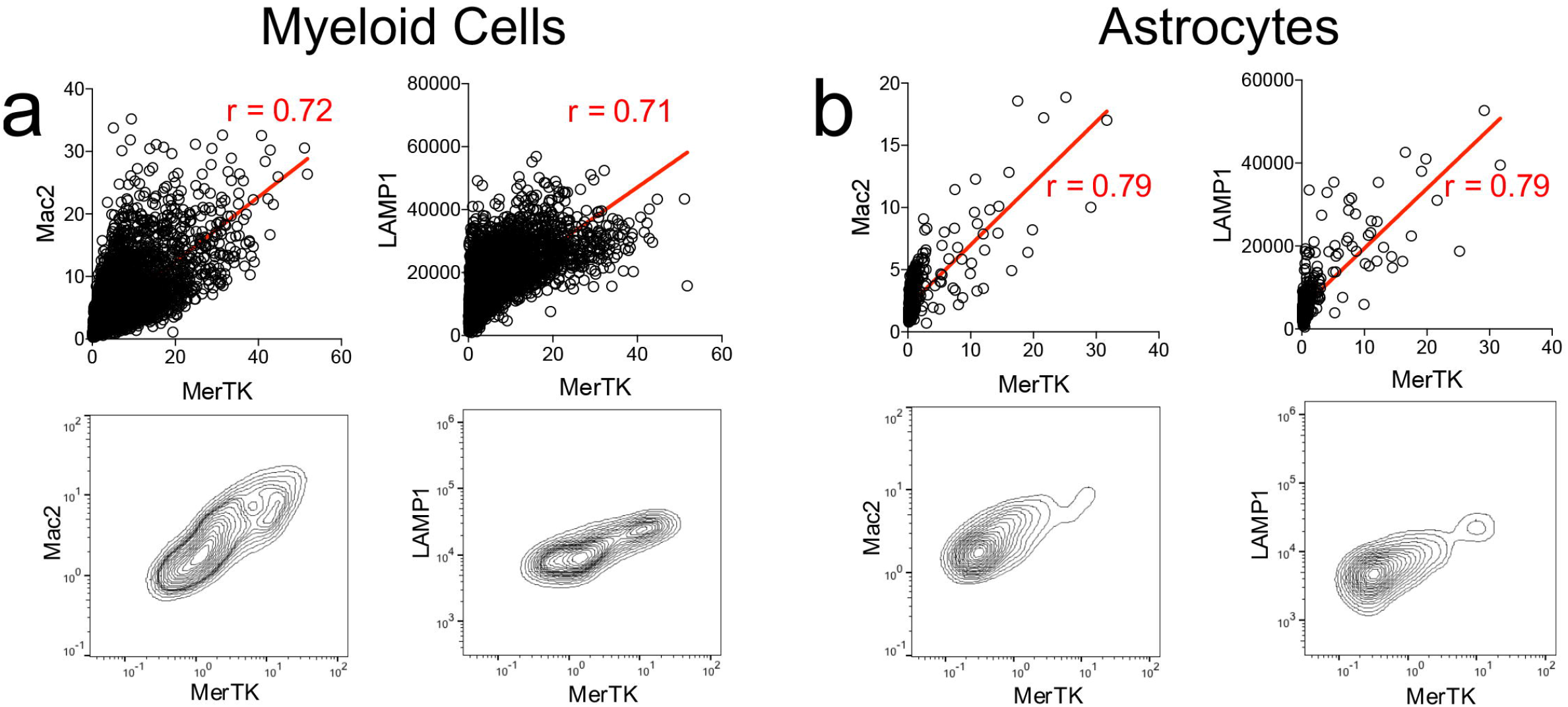
Single-cell marker correlations in the late lesion. **(a, b)** Co-expression of Mac2 and MerTK, and LAMP1 and MerTK in **(a)** myeloid cells (n = 6100), and **(b)** astrocytes (n = 528). Co-expression plots are shown with linear expression values and regression lines with Pearson correlation coefficients, and in flow cytometry contour plot form with log10-transformed expression values

### Phenotypes in the early and late acute lesions engaged in specific cell-cell interactions

We next investigated the spatial relationships between different phenotypes with a computational tool integrated into histoCAT that performs an unbiased, systematic analysis of pairwise phenotype interactions and avoidances [37]. After excluding interactions between cells of the same or spatially adjacent phenotypes, our analysis demonstrated distinct interaction signatures for both lesions, (Fig. 7a, b). At a significance cut-off of p < 0.01, these included interactions between inner rim myeloid phenotype m3 (MerTK and CD45 high) and astrocyte phenotype a1 (Mac2 high), as well as interaction of highly activated myeloid rim phenotype m1 and the rim/center phenotype m4 (HLA-DR and ferritin high) with astrocyte phenotype a2 (HLA-DR high) in the early lesion. In the late lesion, highly activated perivascular macrophages (m4) interacted with most myeloid cell phenotypes and both astrocyte phenotypes. There were also significant interactions between myeloid and astrocyte phenotypes m6 and a1, and among lesion center myeloid phenotypes (m6 with m7 and m8). At a significance cut-off of p < 0.05, we found that T cells in the late lesion interacted with myeloid phenotypes in the perivascular space (m4), which expressed HLA-DR, and in the lesion center (m8).

**Fig. 7.**
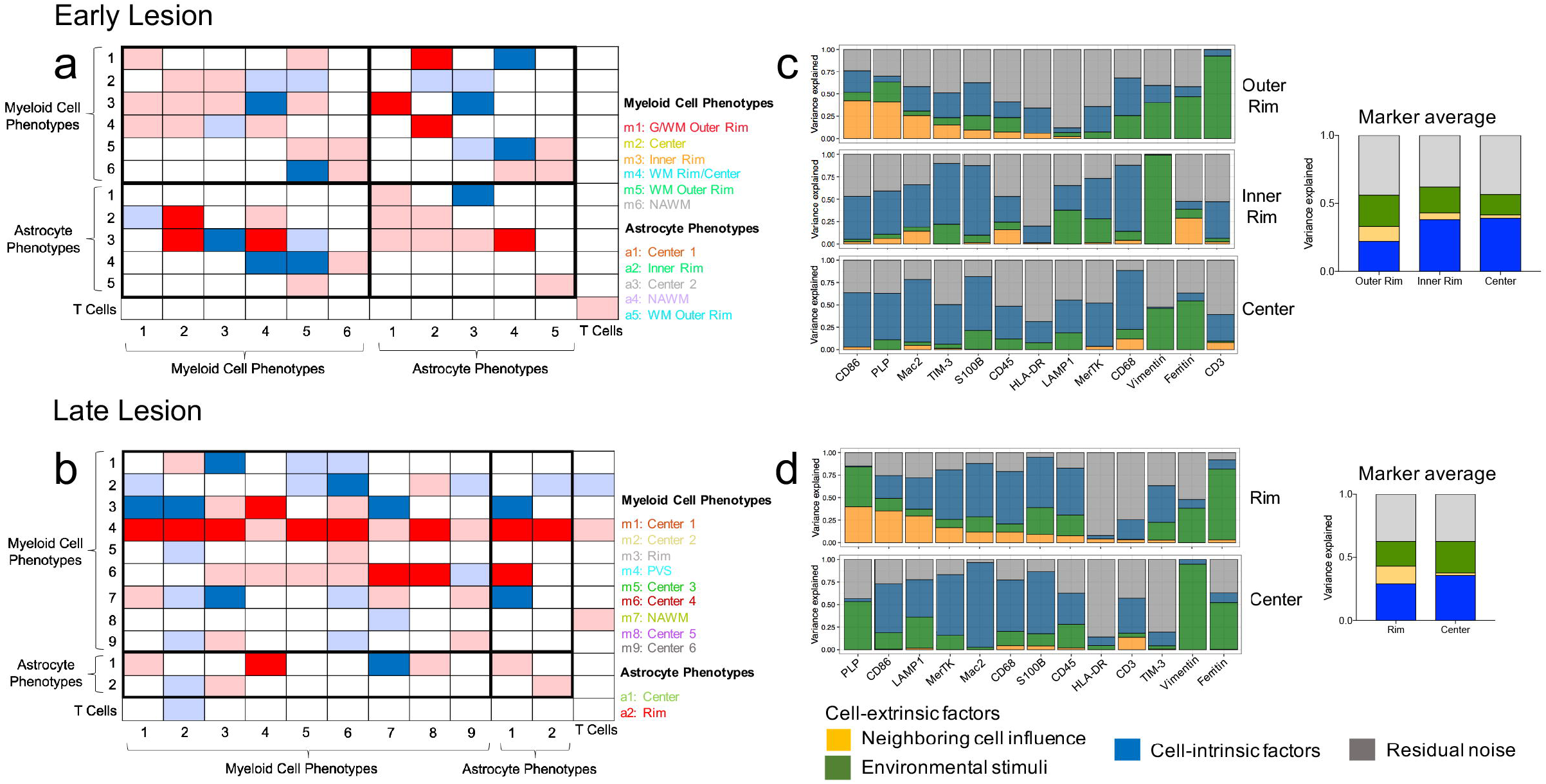
Neighborhood and spatial variance component analyses for the early and late lesions. **(a, b)** Neighborhood analysis heatmaps of all significant pairwise phenotype interactions (red) and avoidances (blue) in the **(a)** early and **(b)** late lesions. White represents no significant spatial relationship. Dark boxes are highly significant spatial relationships (p < 0.01). Lightly shaded boxes are less significant relationships (p < 0.05) and interactions between cells of the same or spatially adjacent phenotypes. Rows visualize the significance of a phenotype surrounded by other phenotypes, and columns visualize the significance of a phenotype surrounding other phenotypes. **(c, d)** Spatial variance component analysis (SVCA) for the **(c)** early and **(d)** late lesions, showing the proportion of marker expression variance attributable to neighboring cell influences, environmental stimuli, cell-intrinsic factors and residual noise in different lesion zones. Additional plots show the average proportion of marker variance attributable to each factor in different lesion zones. G/WM = gray and white matter; WM = white matter; PVS = perivascular space; NAWM = normal-appearing white matter

### Influence of lesion environment on marker expression

Finally, we used spatial variance component analysis (SVCA) to model the effects of extrinsic factors (neighboring cells and non-cellular environmental stimuli) and cell-intrinsic factors on variations in cell marker expression [3]. In the rim of both lesions, the expression of several markers was highly influenced by neighboring cells, including CD86, PLP, and Mac2 in the early lesion, and CD86, PLP and LAMP1 in the late lesion. Other markers, such as ferritin, vimentin, and CD3 (early lesion), and ferritin, vimentin, and TIM-3 (late lesion), were driven predominantly by non-cellular environmental stimuli (Fig. 7c, d). The relative influence of these factors changed toward the inner lesion rim and center, leading to an overall increased impact of cell-intrinsic factors and a decrease in the influence of external factors. In the lesion center, the primary agents influencing marker expression were cell-intrinsic factors and, to a lower degree, environmental stimuli (Fig. 7c, d).

## Discussion

Our study examines the landscape of myeloid and astrocyte phenotypes in early and late acute MS brain lesions using IMC. To our knowledge, this is the first application of highly multiplexed imaging to MS tissue. We applied thirteen markers that are known to be expressed by activated glial cells during MS lesion development. Clustering resulted in eleven myeloid cell and astrocyte phenotypes that localized to distinct lesion areas. Moreover, individual phenotypes interacted selectively with other cell types, and marker expression was driven by different factors in cells within the lesion rim than the center. Thus, our approach provides a wealth of data on cellular spatial organization that is not accessible with standard histology.

The alignment of myeloid cell phenotypes with different lesional layers suggests functional specificity and validates our clustering approach. This spatial separation was most pronounced in the early lesion, but was lost in the center of the late lesion, where phenotypes were intermixed. In addition, marker expression was the highest in myeloid phenotypes located at the lesion rim and diminished substantially towards the lesion center in both lesions. Consistent with the different stages of myelin phagocytosis and degradation, the myeloid phenotypes in the rim were larger than those in the lesion center. An additional feature of the late lesion was the presence of numerous, highly activated macrophages in perivascular spaces throughout the lesion. Since these macrophages are believed to transition into the vasculature [22], this may indicate they exit the CNS in a highly activated state. In contrast to myeloid cells, marker expression in astrocyte phenotypes did not follow a rim-to-center gradient, but was comparable throughout the lesion.

Using PHATE mapping and Pseudotime to determine transitions between phenotypes, we found that myeloid cell or astrocyte phenotypes did not develop along a linear transition continuum, e.g. from the outer to the inner lesion rim and lesion center. Instead, the different phenotypes branched into multiple independent trajectories, suggesting that they emerge within the lesion environment independent from each other. We cannot exclude that the parametric depth of our study is too low to detect transition states, especially since Pseudotime has been developed to delineate phenotypic trajectories from whole transcriptome sequencing data [41].

Neighborhood analysis demonstrated distinct cellular interaction signatures for both lesions, e.g. between phagocytic inner rim macrophages and center astrocytes in the early lesion, and between T cells and two myeloid phenotypes in the late lesion. This indicates that cellular interactions are not random in this hypercellular lesion environment, but occur between specific subpopulations and cell types such as lymphocytes. The low parametric depth of our study does not allow us to identify the functional implications of these interactions; however, they may represent nodal points of cellular communication critical for lesion formation and maintenance of low-grade inflammation.

Finally, spatial variance component analysis (SVCA) suggests that cell-extrinsic factors drive marker expression to a higher degree in the lesion rim than in the center. Conversely, cell-intrinsic factors have a more prominent influence on marker expression in the lesion center. This suggests that glia cells in the lesion rim respond more to cues from the microenvironment, such as cytokines or receptor-ligand interactions, while glial activation in the lesion center is the result of cell-intrinsic programs such as cellular changes set in motion by myelin phagocytosis.

Myeloid cell/microglial heterogeneity has recently been examined by us and others with single-cell RNA sequencing in healthy CNS, MS lesions and neurological diseases such as Alzheimer’s disease, Parkinson’s disease and temporal lobe epilepsy [23,28]. In keeping with our results, these efforts have identified multiple myeloid cell/microglial phenotypes; specifically, Masuda and colleagues determined nine myeloid cell phenotypes in MS lesions and healthy brain [23], and we found ten myeloid phenotypes across multiple neurodegenerative diseases [28]. In the latter study, we identified one microglia cluster (cluster 7), characterized by high expression of CD74, which was enriched for genes associated with MS susceptibility. This cluster was the most congruent with our rim phenotypes, as it was enriched for several genes that were highly expressed at the rim of our lesions, i.e. LAMP1, CD45, CD68, CD86 and HLA-DR. Moreover, we confirmed that this cluster localized to the lesion by staining our lesions for CD74 (data not shown). Other attempts to cluster myeloid cells in experimental autoimmune encephalomyelitis (EAE), a mouse model of MS, using single-cell cytometry [26], and in MS lesions using single nuclear RNA sequencing [14], have yielded substantially less myeloid cell heterogeneity.

Our study is limited by the small sample size and the low number of markers, which may result in inaccurate phenotype clustering. However, as a proof-of-concept study, it demonstrates the ability of multiplexed tissue imaging and appropriate single-cell analytics to reveal the heterogeneity and spatial properties of glial cell phenotypes in MS lesions.

## Conclusions

In summary, we found that phenotypic clustering based on differential expression of thirteen glial activation markers produced multiple myeloid cell and astrocyte phenotypes that occupied specific lesion zones. Myeloid cells were activated along a rim-to-center axis, and specific myeloid cell-astrocyte-lymphocyte interactions were present in both lesions. Our study highlights the potential of imaging mass cytometry, paired with novel computational tools, to provide insight into lesion-forming phenotypes and their spatial organization in MS lesions.

In ongoing studies, we are combining cell clustering based on single-nuclear RNA sequencing data of MS lesion-derived glial cells with highly multiplexed imaging to simultaneously obtain maximal parametric depth and high spatial resolution at single-cell level. The results will help define the phenotypes and key interaction networks that drive active demyelination and chronic, low-grade inflammation in established lesions and may ultimately provide targets for therapeutic intervention in relapsing and progressive MS.

## List of abbreviations

MS: multiple sclerosis
CNS: central nervous system
IMC: imaging mass cytometry
CyTOF: mass cytometry
MBP: myelin basic protein
DAB: 3,3-diaminobenzidene
PHATE: Potential of Heat-diffusion Affinity-based Transition Embedding
SVCA: spatial variance component analysis
WM: white matter
G/WM: gray and white matter
NAWM: normal-appearing white matter
G/WMoR: gray and white matter outer rim
WMoR: white matter outer rim
iR: inner rim
WM R/C: white matter rim/center
C: center
R: rim
PVS: perivascular space
EAE: experimental autoimmune encephalomyelitis

## Additional file

Additional file 1 (DOCX): Tables S1-S2 and Figures S1-S8.

## Declarations

### Ethics approval and consent to participate

Full ethical approval for the use of de-identified, human CNS autopsy tissue was obtained according to Institutional Review Board-approved protocols (Yale Human Investigation Committee), and informed consent was received from all human participants.

### Consent for publication

Not applicable.

### Availability of data and materials

The datasets generated and/or analyzed during the current study as well as R code are available in a GitHub repository, https://github.com/PittLab/IMC_Park_et_al_2019.

### Authors’ contributions

D.P., C.P. and G.P. conceived and designed the study; C.P. performed the experiments; C.P., G.P. and D.P. analyzed the data; G.P., M.L.-R., E.B., E.C.S., P.L.D., B.M.S. and D.P. provided methodological, technical and/or scientific assistance; E.C.S. provided IMC tissue imaging service; C.P., G.P., M.L.-R., E.C.S., P.L.D., B.M.S. and D.P. reviewed the manuscript; C.P. and D.P. wrote the manuscript.

## Acknowledgments

The authors thank Fluidigm Corporation (Markham, ON, Canada) and the Yale University CyTOF core facility for providing IMC tissue imaging service and metal-conjugated antibodies, as well as Denis Schapiro, Damien Arnol and Kevin Moon for expert assistance with computational analyses.

