## Additional file 1 for "The landscape of myeloid and astrocyte phenotypes in acute multiple sclerosis lesions"

**CONTENTS**

**Supplementary Tables	Page**

Table S1 (Antibodies used for brightfield histology)	2

Table S2 (Metal-conjugated antibodies used for IMC)	3

**Supplementary Figures**

Figure S1	4

Figure S2	5

Figure S3	6

Figure S4	7

Figure S5	8

Figure S6	9

Figure S7	10

Figure S8	11

**Additional file 1: Table S1** Antibodies used for brightfield histology

| **Primary Antibody** | **Host** | **Clone** | **Company** | **Target/Significance** | **Primary Antibody Dilution** | **Secondary Antibody** | **Secondary Antibody Dilution** |
| --- | --- | --- | --- | --- | --- | --- | --- |
| MBP | Rat | 12 | Millipore-Sigma | Myelin Basic Protein | 1:500 | Invitrogen, 629540 | 1:500 |
| CD68 | Rabbit | D4B9C | Cell Signaling | Activated macrophage/microglia transmembrane glycoprotein | 1:500 | Vector Laboratories, BA-1000 | 1:500 |
| MAP2 | Mouse | 5H11 | Novus | Neuron structural protein | 1:200 | Vector Laboratories, BA-9200 | 1:500 |

| **Antibody** | **Clone** | **Company** | **Target/Significance** | **Conjugated Metal Isotope** | **Conjugation Source** | **Dilution** |
| --- | --- | --- | --- | --- | --- | --- |
| LAMP1/CD107a | H4A3 | Fluidigm | Lysosomes | 151Eu | Fluidigm | 1:100 |
| CD3 | Polyclonal | Fluidigm | T cells | 170Er | Fluidigm | 1:75 |
| CD45 | 2B11 | Fluidigm | Leukocyte activation | 152Sm | Fluidigm | 1:100 |
| CD68 | KP1 | Fluidigm | Activated macrophages/microglia | 159Tb | Fluidigm | 1:300 |
| CD86 | Polyclonal | R&D | “M1” co-stimulatory molecule | 171Yb | Our Lab | 1:100 |
| Ferritin heavy chain | 1-2.3.1.2 | Millipore-Sigma | Iron storage | 158Gd | Our Lab | 1:400 |
| HLA-DR | YE2/36 HLK | Fluidigm | MHC Class II | 174Yb | Fluidigm | 1:200 |
| Mac2 | M3/38 | Fluidigm | “M2” activation | 153Eu | Fluidigm | 1:600 |
| MerTK | y323 | abcam | “M2” phagocytosis | 148Nd | Our Lab | 1:50 |
| PLP | plpc1 | Bio-Rad | Myelin | 142Nd | Our Lab | 1:100 |
| S100B | S100B/1706R | Novus | Astrocytes | 167Er | Our Lab | 1:400 |
| TIM-3 | D5D5R | Fluidigm | Co-inhibitory molecule | 154Sm | Fluidigm | 1:100 |
| Vimentin | RV202 | Fluidigm | Hypertrophic astrocytes | 143Nd | Fluidigm | 1:100 |

**Additional file 1: Table S2** Metal-conjugated antibodies used for IMC

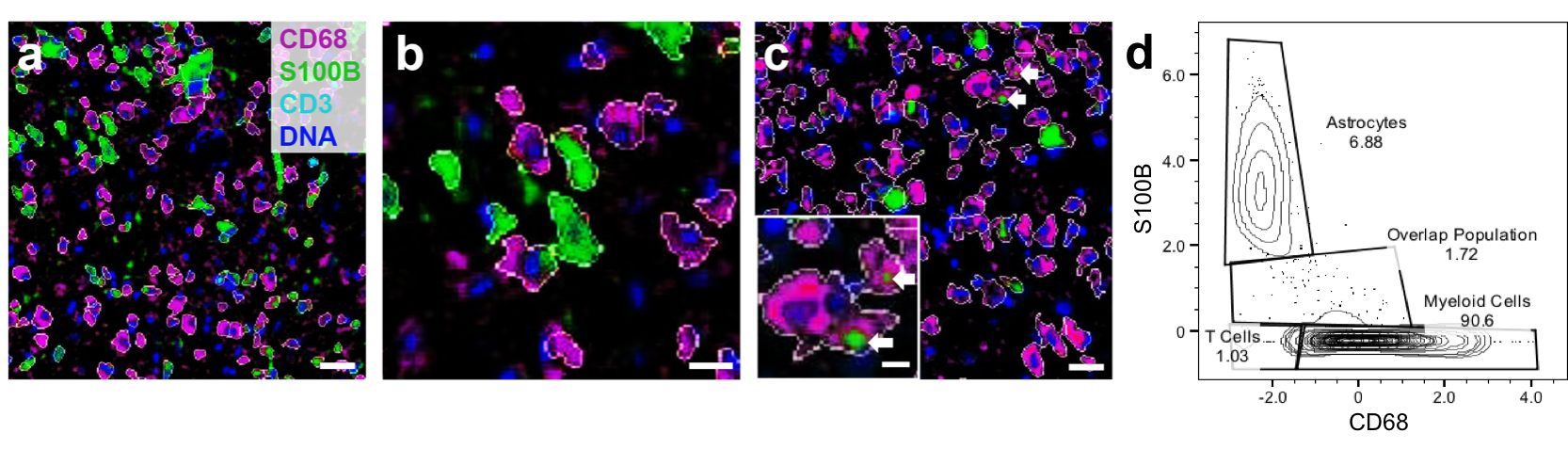
**Additional file 1: Figure S1** Single-cell segmentation. **(a-c)** Segmentation of myeloid cells (CD68, magenta), astrocytes (S100B, green) and T cells (CD3, cyan) performed on CellProfiler. **(b)** shows cells in **(a)** at higher magnification. **(c)** shows examples of S100B^+^ astrocyte processes (white arrows) that closely neighbor CD68^+^ cells, which resulted in the co-segmentation of these markers in a small fraction of cells. **(d)** Gating of CD68^+^ and S100B^+^ populations on a flow cytometry plot as a quality control for segmentation (late lesion). The overlap population consists of cells with co-segmented CD68 and S100B like those in **(c)**. Scale bar in **(a)** = 30 μm; **(b)** = 15 μm; **(c)** = 30 μm, inset = 10 μm. Z-score normalized expression intensities are shown **(d)**

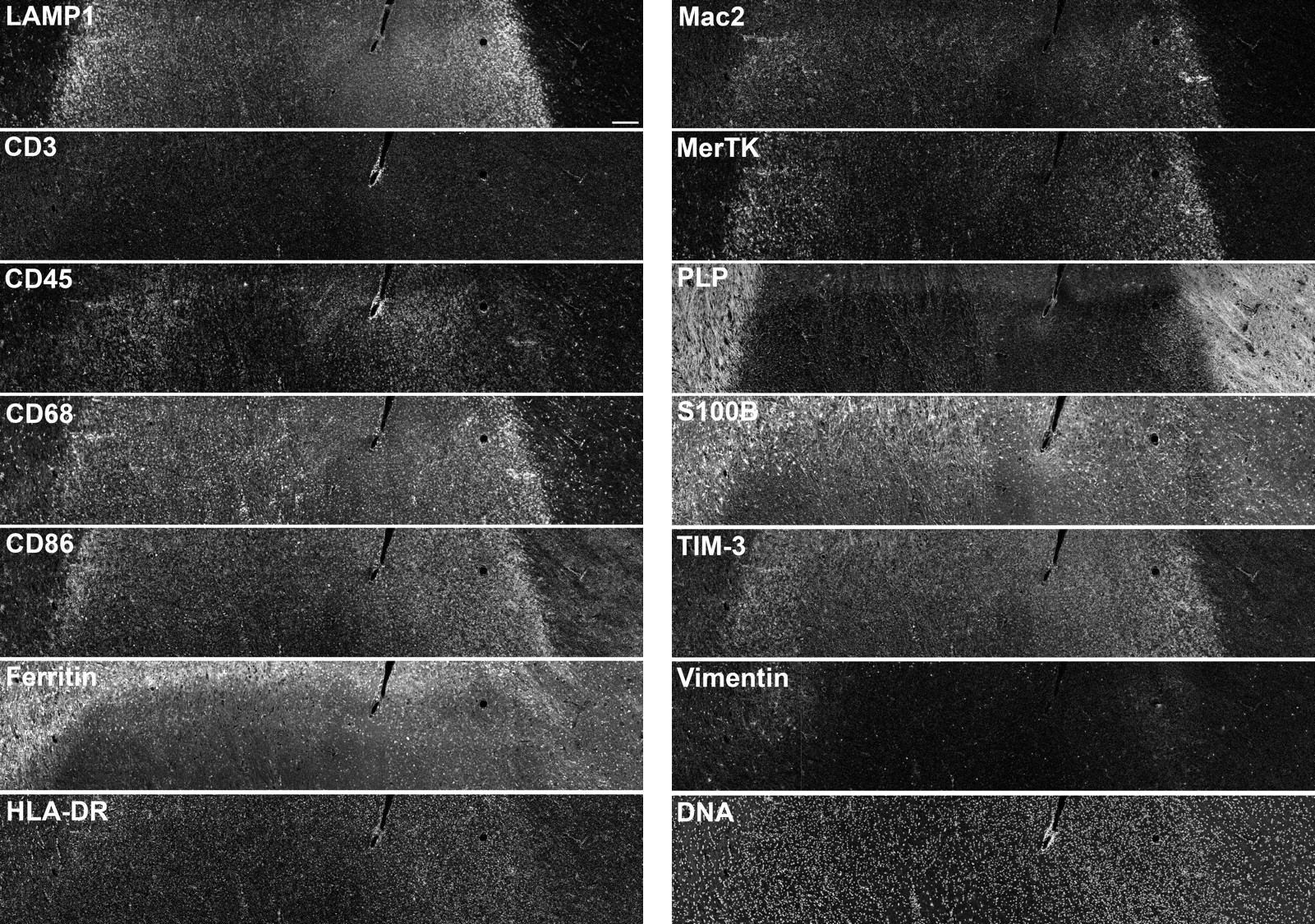

**Additional file 1: Figure S2** Imaging mass cytometry histology images of the early lesion. The thirteen markers used for cell clustering as well as DNA counterstaining are shown. Scale bar = 200 μm

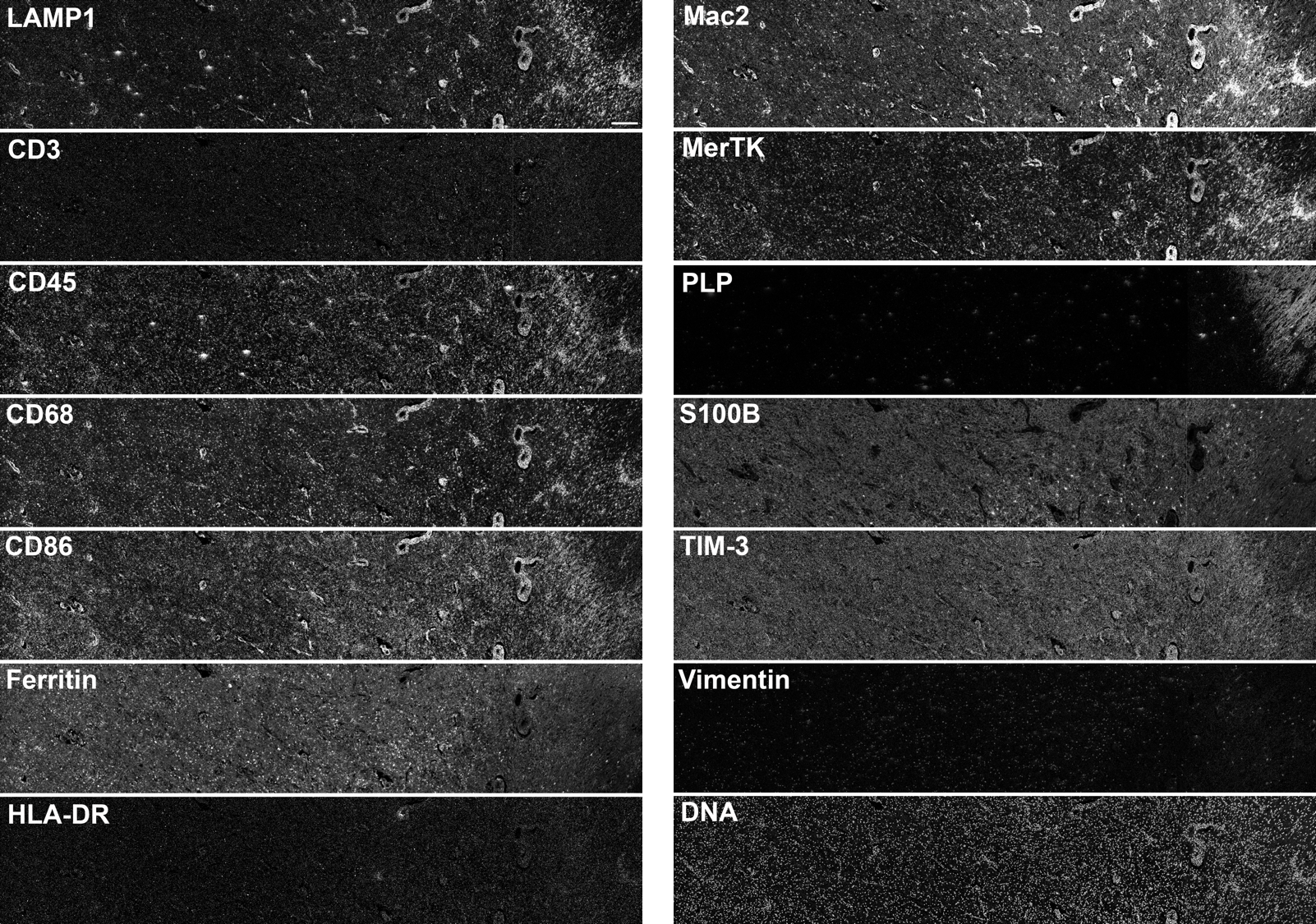

**Additional file 1: Figure S3** Imaging mass cytometry histology images of the late lesion. The thirteen markers used for cell clustering as well as DNA counterstaining are shown. Scale bar = 200 μm

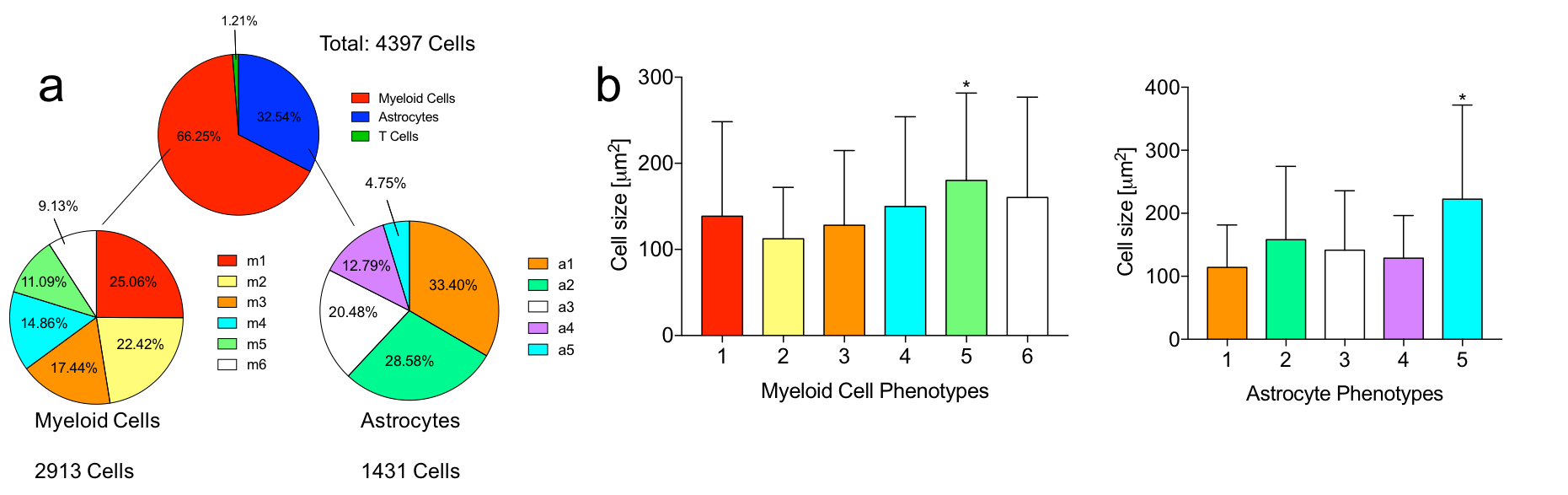

**Additional file 1: Figure S4** Early lesion cell phenotype frequency and size. **(a)** Relative quantities of each cell phenotype. **(b)** Cell phenotype size. Cells of myeloid phenotype 5 (m5), which occupy the white matter outer rim, are significantly larger than all other myeloid phenotypes. Similarly, cells of astrocyte phenotype 5 (a5), which occupy the white matter outer rim, are significantly larger than all other astrocyte phenotypes. Data represent means + standard deviation. *p < 0.0001 by one-way ANOVA followed by the Tukey-Kramer multiple comparison test

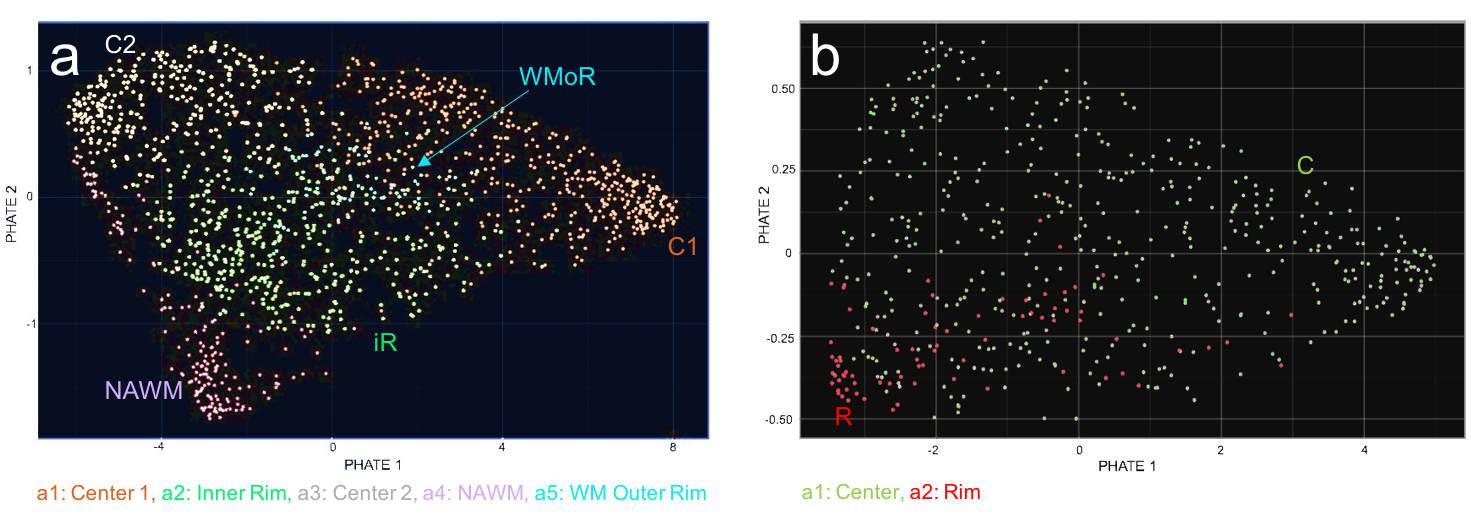

**Additional file 1: Figure S5** PHATE mapping of astrocyte phenotypes in the **(a)** early and **(b)** late lesion, demonstrating no linear phenotype transition in either lesion. R = rim; WMoR = white matter outer rim; iR = inner rim; C = center; NAWM = normal-appearing white matter

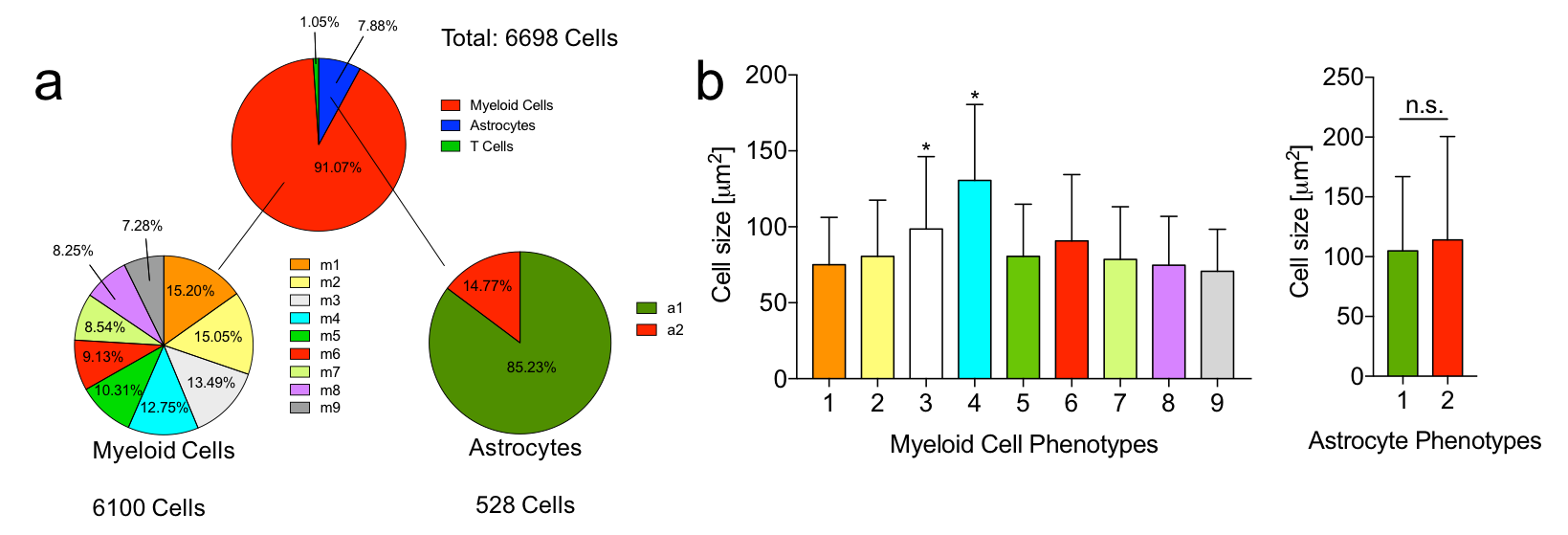

**Additional file 1: Figure S6** Late lesion cell phenotype frequency and size. **(a)** Relative quantities of each cell phenotype. **(b)** Cell phenotype size. Cells of myeloid phenotypes 3 and 4 (m3, m4), in the rim and perivascular space, respectively, are significantly larger than all other myeloid phenotypes, which occupy the lesion center. Data represent means + standard deviation. *p < 0.0001 by one-way ANOVA followed by the Tukey-Kramer multiple comparison test. n.s. = not significant

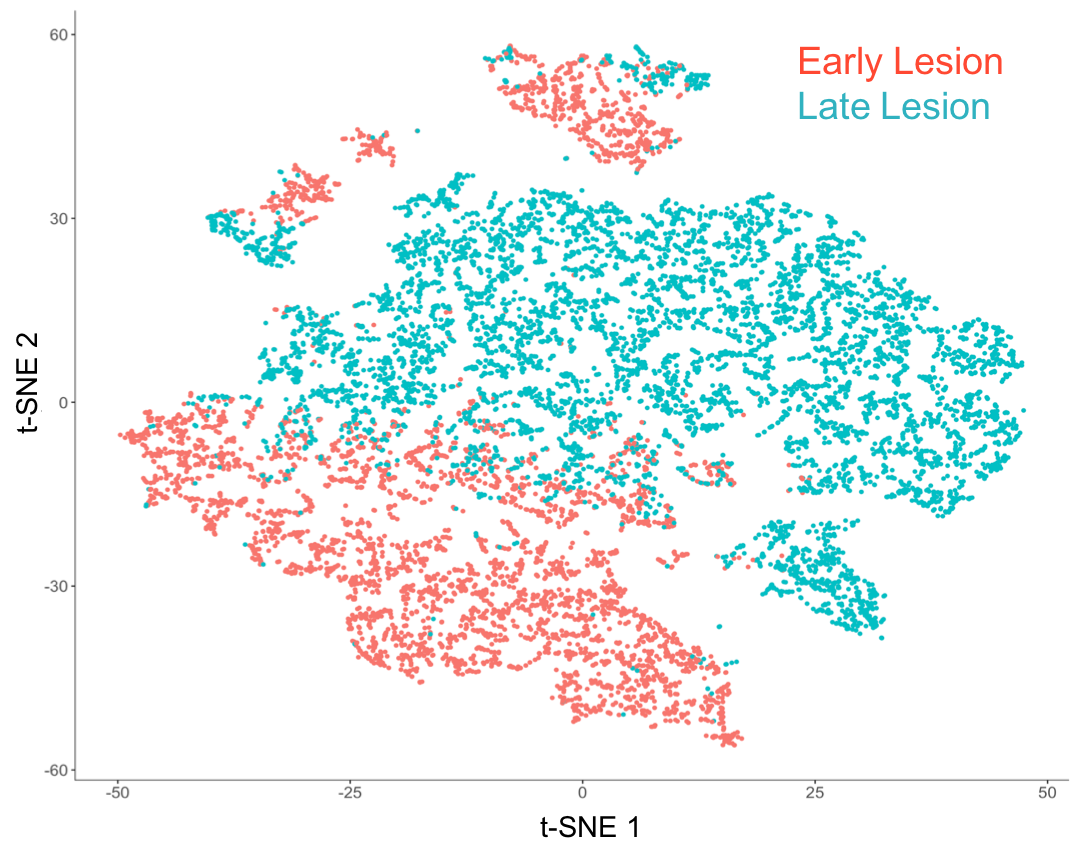

**Additional file 1: Figure S7** Comparison of lesion cell character with t-SNE. The plot shows all analyzed cells from the early and late lesions. Cellular populations overlap to some degree but are predominantly distinct

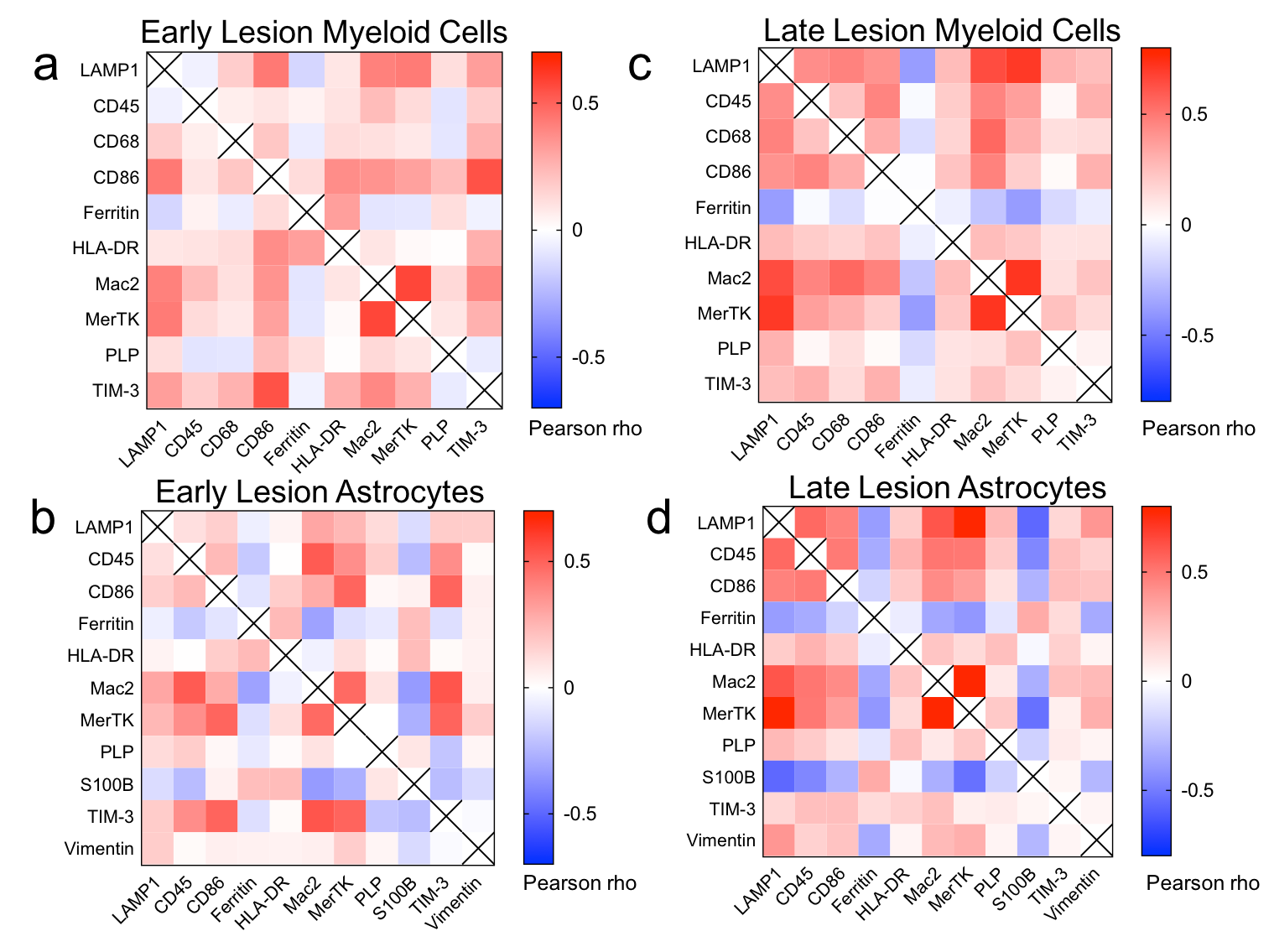

**Additional file 1: Figure S8** Pearson correlation matrices for myeloid cells and astrocytes. Matrices for early lesion **(a)** myeloid cells (n = 2913) and **(b)** astrocytes (n = 1431), and late lesion **(c)** myeloid cells (n = 6100) and **(d)** astrocytes (n = 528) are shown
